## Supplemental Information for "Dissecting reversible and irreversible single cell state transitions from gene regulatory networks"

### Supplemental Materials

### Supplementary Figures

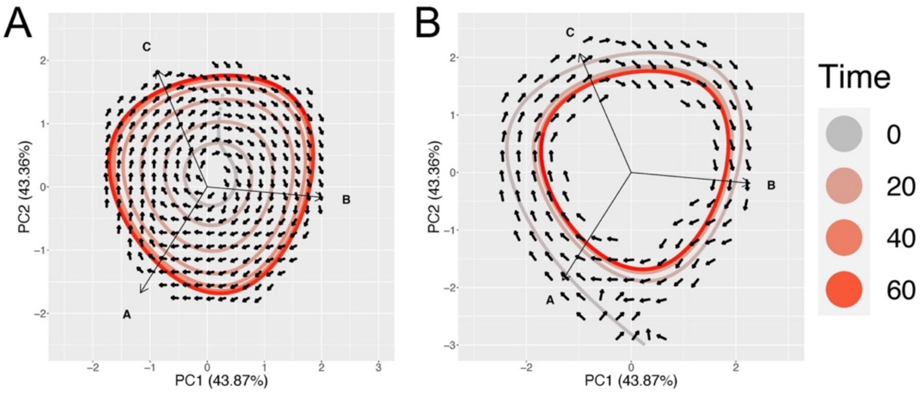


**Supplementary Figure 1.** Repressilator (REP) time trajectory simulations. STICCC predictions (vectors shown as arrows) using the gene expression snapshots from the simulated REP trajectories approaching the limit cycle (with later time points indicated in red) from two different initial conditions (with earlier time points indicated in gray). The plots show the projection of the time-series gene expression data onto the first two principal components from the simulated gene expression of an ensemble of 10000 models.

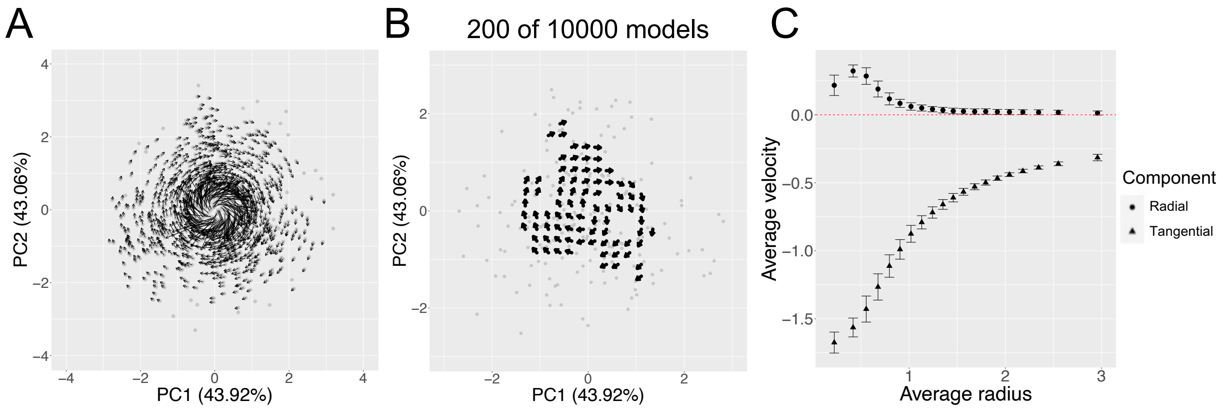


**Supplementary Figure 2.** Repressilator cell-specific vectors and down-sampling. **A)** Cell-specific outgoing transition vectors $v_1$ calculated for a simulated gene expression of an ensemble of 10000 REP models. Shown is a random subset of 1000 cells and their predicted vectors. **B)** State transition pattern is conserved even with extreme under-sampling. Grid-smoothed vector field for REP circuit is shown after calculating vectors on a subset of 200 out of 10000 models. **C)** The panel shows the mean values of the radial (circles) and tangential (triangles) components of the inferred vectors for cells in various radial bands around the origin of the gene expression space, projected onto the first two principal components. The inferred transition vectors are predominantly dominated by the tangential components, suggesting oscillatory state transitions. Error bars are drawn at +/- 1 standard deviation.

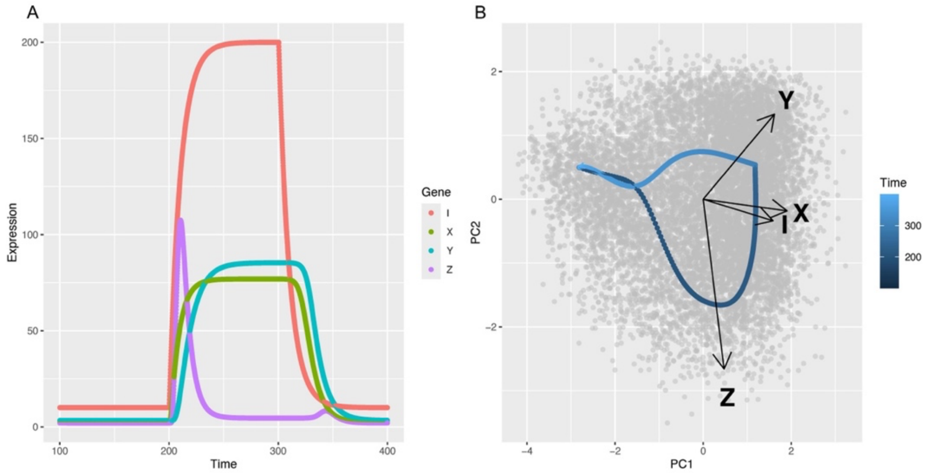

**Supplementary Figure 3.** Time trajectory for incoherent feedforward loop (FFL) simulation. A model with parameters as described in **Table S1** was simulated deterministically with an increase in production rate for signal node I at t=200, which was then removed at t=300. **A)** Line plot of expression values over time for each gene in the network, colored by gene. **B)** Time trajectory from (A) projected to the first two principal components from ensemble simulation of the FFL circuit, with ensemble data shown in grey and the trajectory plotted on top, colored by time (dark blue to light blue). Loading vectors for each gene in the first two principal components are also plotted and labeled with arrows.

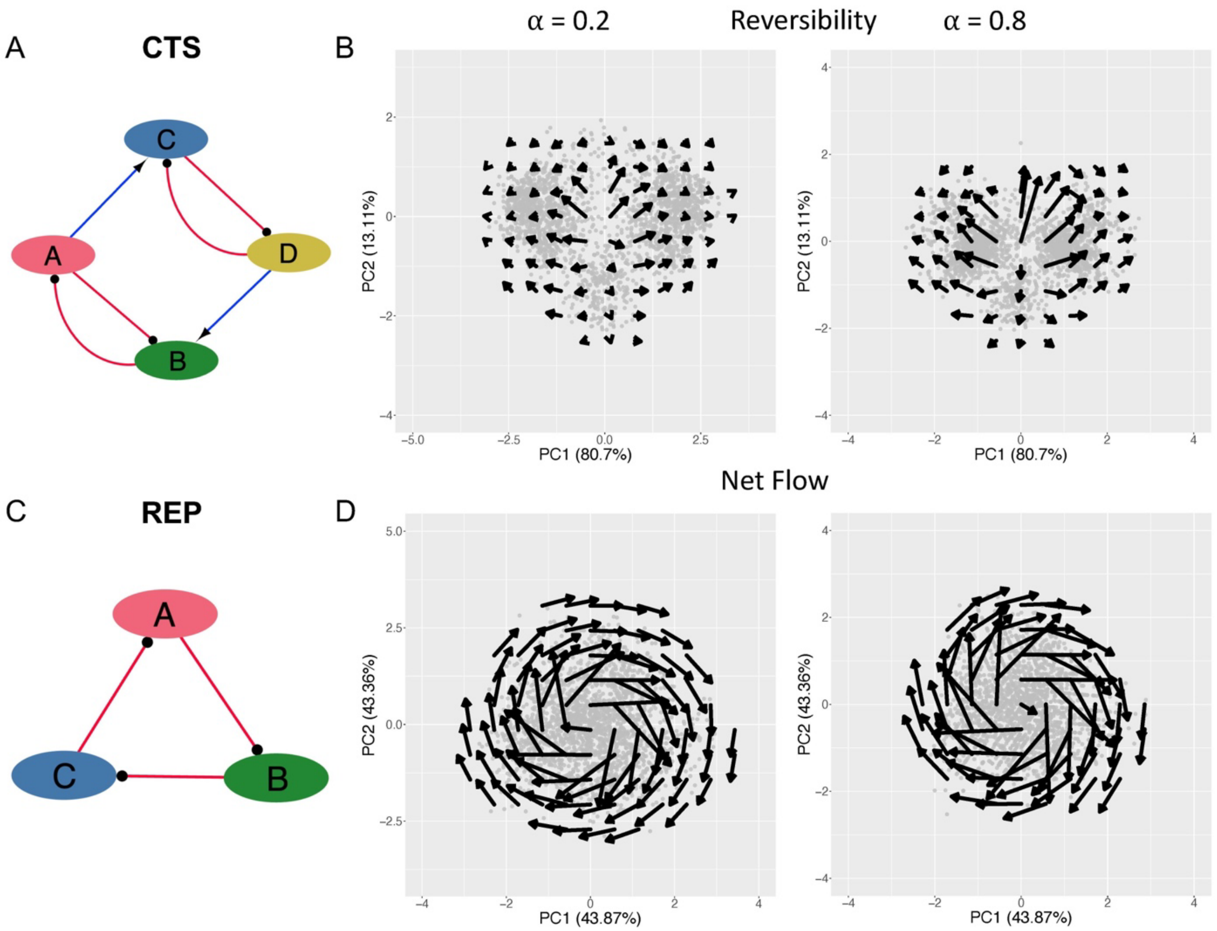


**Supplementary Figure 4.** Vector fields for coupled toggle switch (CTS) and repressilator circuits at low and high levels of simulated noise. **A)** CTS topology diagram. **B)** Reversibility vector field for CTS at low (a = 0.2) and high (a = 0.8) dropout levels. **C)** Repressilator topology diagram. **D)** Net flow vector field for repressilator at low and high dropout levels.

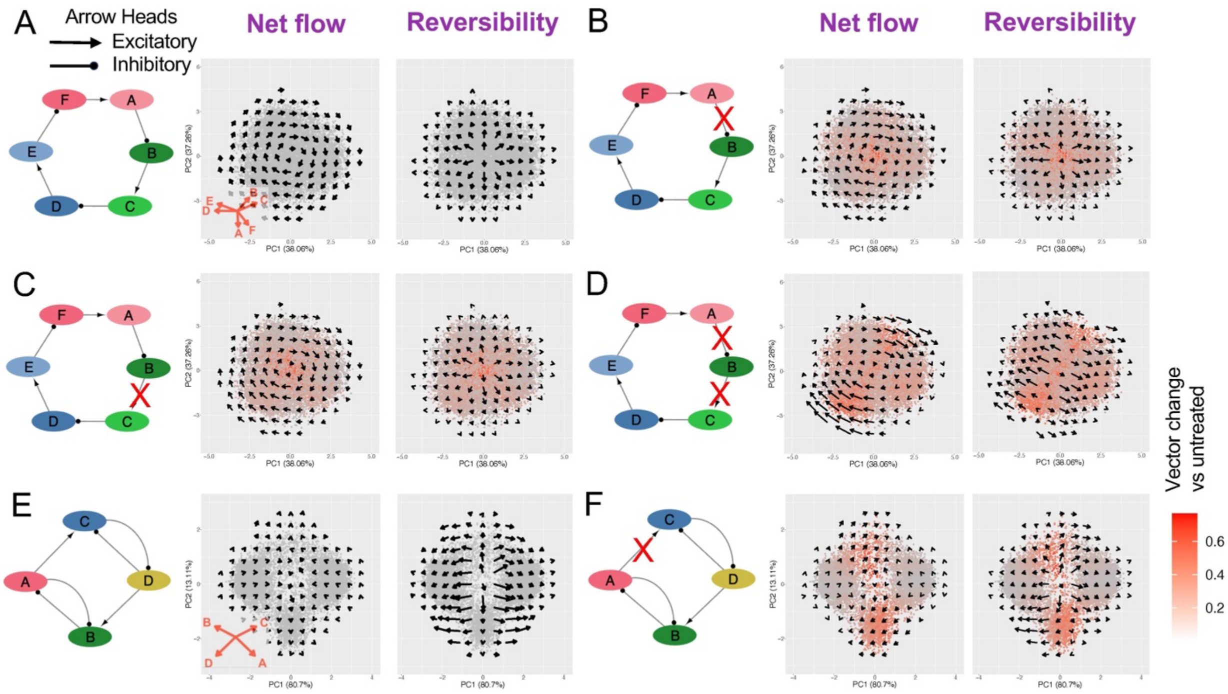


**Supplementary Figure 5.** Edge sensitivity analysis reveals roles and importance of network edges. Using simulations of a 6-gene repressilator and CTS, different subsets of the true topology were provided to STICCC to infer edge importance based on the resulting change in vectors. Point color denotes relative change in vector prediction. **A)** Expanded repressilator with vector predictions based on the full topology projected on PCA. **B-D)** Predictions for the expanded repressilator with one or two edges omitted. **E-F)** Predictions for CTS with full topology and the edge from A to C omitted.

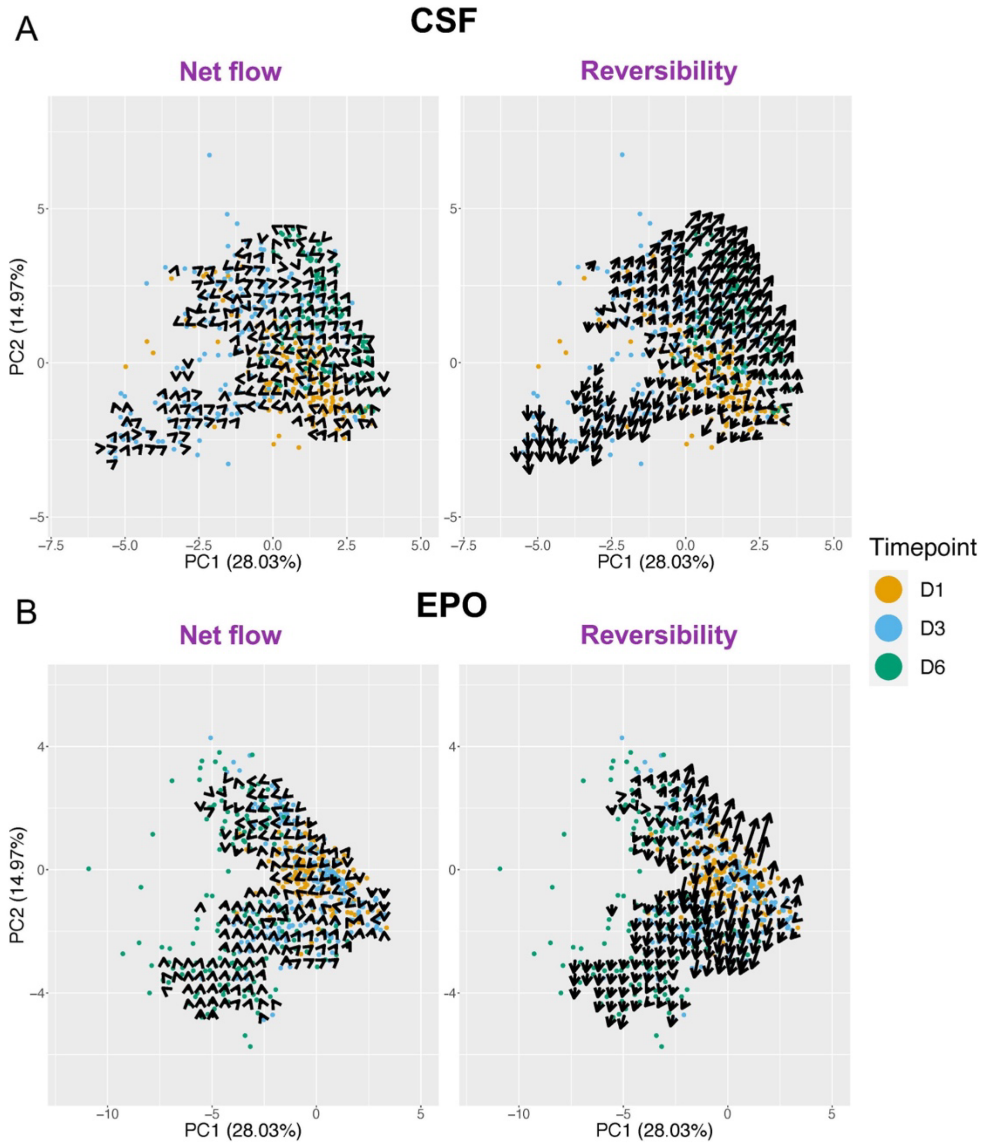


**Supplementary Figure 6.** STICCC predictions for hematopoietic stem cell data separated by treatment condition. Point color denotes timepoint. **A)** Net flow and reversibility predictions for HSC cells treated with CSF. **B)** Predictions for HSC cells treated with EPO.

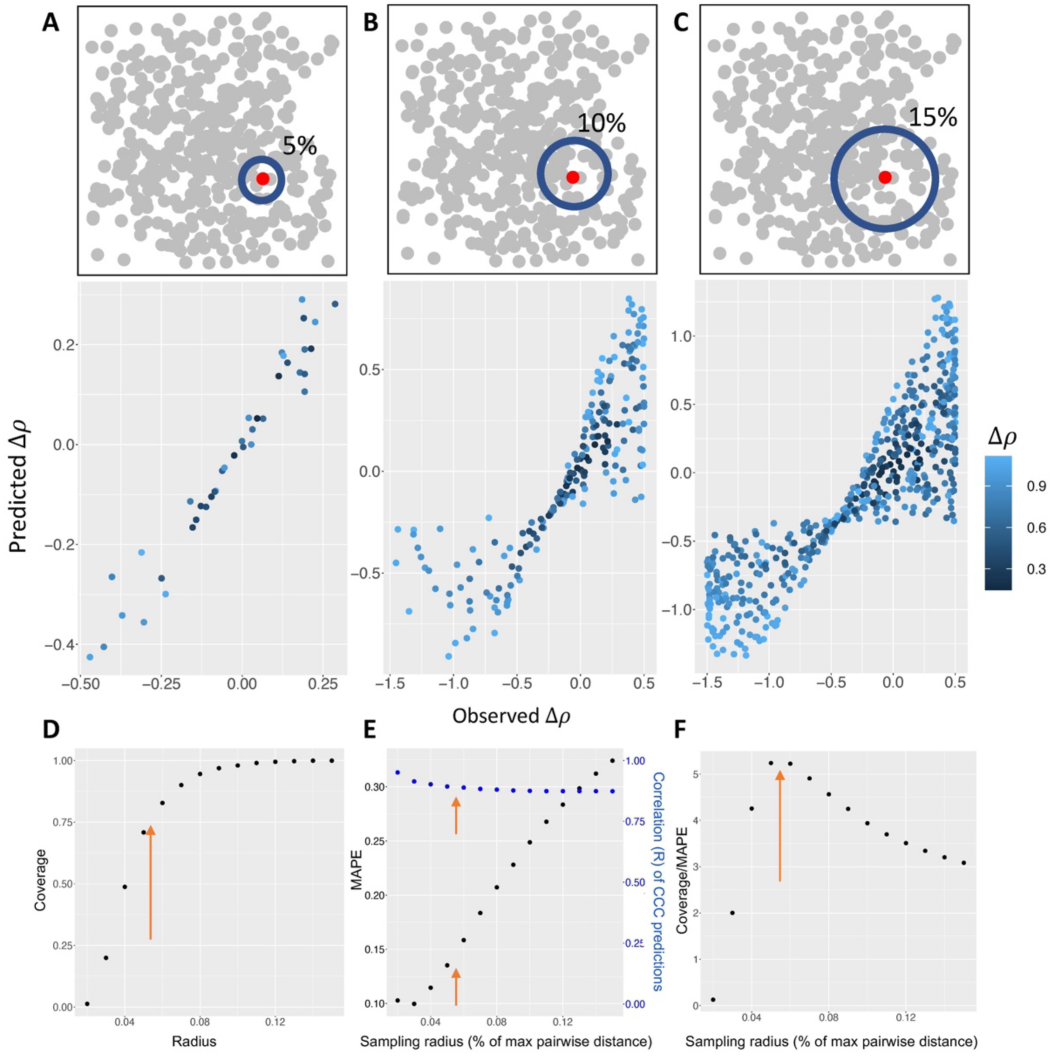


**Supplementary Figure 7.** Linear regression quality depends on sampling radius. A-C) Example neighborhoods for a sampling radius of 5%, 10%, and 15% of the maximum pairwise Euclidean distance in gene expression between cells, respectively, illustrated by a blue circle around a red center cell. Below, scatterplots show observed and predicted CCC values for neighboring cells. D) Coverage, *i.e.*, proportion of cells for which a prediction is generated, as a function of sampling radius. E) Median absolute percent error (MAPE), left axis, and R value of linear regression, right axis, for various sampling radii. F) Ratio of coverage to MAPE for various sampling radii.

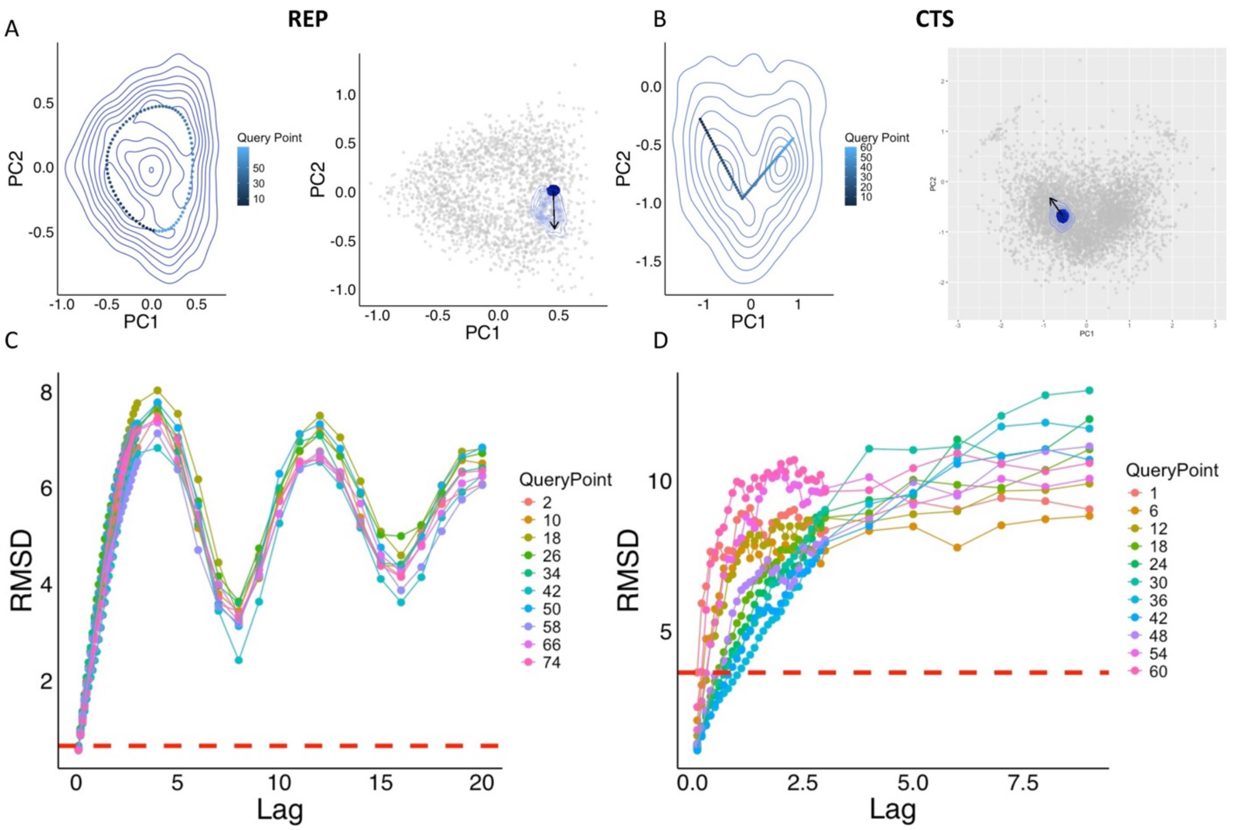


**Supplementary Figure 8.** Selection of lag periods for time trajectory comparisons. **A)** Left panel: Density plot of gene expression snapshots from a noisy time trajectory simulation of the REP circuit with blue contour lines, projected onto the first two principal components from the simulated gene expression of an ensemble of 10000 models. Points shown are snapshots of the deterministic limit cycle, colored and indexed 1-77 with earlier timepoints in black and later timepoints in blue. Right panel: Illustrative example of points selected to compare with predicted angles. Points in dark blue are sampled from the noisy trajectory by proximity to a selected point from the deterministic cycle. Blue contour lines show the time evolution of each point in the previous group after a time lag of 0.5. Arrow shows the predicted $v_1$ vector for the selected point from the limit cycle. **B)** Left panel: Density plot of gene expression snapshots from a noisy time trajectory simulation of CTS circuit with blue contour lines, projected onto the first two principal components from the simulated gene expression of an ensemble of 10000 models. Points shown are a linear interpolation between the medians of three clusters identified with k-means clustering, colored and indexed 1-60 with earlier timepoints in black and later timepoints in blue. Right panel: Illustrative example of points selected to compare with predicted angles. Points in dark blue are sampled from the noisy trajectory by proximity to a selected point from the linear path. Blue contour lines show the time evolution of each point in the previous group after a time lag of 0.5. Arrow shows the predicted $v_2$ vector for the selected point from the linear path. **C)** Line plot of root mean square deviation (RMSD) between initial timepoints and final timepoints by lag period, for 10 points from the REP deterministic limit cycle (indicated by color). Target RMSD, selected based on the highest RMSD of any query point at a lag of 0.1, is indicated by a red horizontal dashed line. **D)** RMSD by lag time between initial and final timepoints for 11 points from the CTS linear transition path, indicated by color. Target RMSD is selected and drawn as in (C). Lag times for REP and CTS were selected independently for each point based on target RMSD; as such, the lag time for the analysis of REP in Fig. 3D was generally constant, whereas for CTS the lag time varies based on the position of the initial states.

### Supplementary Tables

**Supplementary Table 1.** Parameters for the incoherent feedforward loop trajectory simulation.

| Parameter | Gene/Edge | Value |
|---|---|---|
| **Signal Production Rate, G** | I (off) | 1 |
| | I (on) | 20 |
| | X | 10 |
| | Y | 10 |
| | Z | 30 |
| **Degradation Rate, k** | I | 0.1 |
| | X | 0.13 |
| | Y | 0.1 |
| | Z | 0.15 |
| **Hill Coefficient, n** | All | 4 |
| **Fold Change, $\lambda$** | I activates X | 50 |
| | X activates Y | 30 |
| | Y inhibits Z | 50 |
| | X activates Z | 100 |
| **Threshold, $X_0$** | I activates X | 30 |
| | X activates Y | 50 |
| | Y inhibits Z | 20 |
| | X activates Z | 20 |
| **Simulation Time** | - | 400 |

Simulations were conducted with the signal node I in the ‘off’ state for 200 time units, followed by 100 time units with the signal in the ‘on’ state, and a final 100 in the original state.

**Supplementary Table 2.** Parameters for stochastic simulation of the repressilator circuit.

| Parameter | Value |
|---|---|
| Production Rate, G | 30 |
| Degradation Rate, k | 0.5 |
| Hill Coefficient, n | 4 |
| Fold Change, $\lambda$ | 50 |
| Threshold, $X_0$ | 20 |
| Simulation Time | 10000 |
| Simulation Noise, $\xi$ | 0.1 |

**Supplementary Table 3.** Parameters for stochastic simulation of the CTS circuit.

| Parameter | Value |
|---|---|
| Production Rate, G | 50 |
| Degradation Rate, k | 0.1 |
| Hill Coefficient, n | 4 |
| Fold Change, $\lambda$ | 10 |
| Threshold, $X_0$ | 100 |
| Threshold (modified), $X_{0M}$ | 80 |
| Simulation Time | 100000 |
| Simulation Noise, $\xi$ | 2 |

Most parameters are constant for all genes and edges, except a modified threshold value for the edges from B to A and C to D, which increases the activity of these two edges.

**Supplementary Table 4.** Genes used as measurements for cell cycle network nodes.

| Systematic Name | Standard Name | Network Node Name |
|---|---|---|
| YAL040C | CLN3 | Cln3 |
| YAR007C | RFA1 | MBF |
| YIL066C | RNR3 | Whi5 |
| YER111C | SWI4 | SBF |
| YMR199W | CLN1 | Cln1,2 |
| YGR109C | CLB6 | Clb5,6 |
| YLR079W | SIC1 | Sic1 |
| YGL003C | CDH1 | Cdh1 |
| YDR225W | HTA1 | DNA Synthesis (DNA-S) |
| YGR108W | CLB1 | Clb1,2 |
| YMR043W | MCM1 | Mcm1 |
| YGL116W | CDC20 | Cdc20 |
| YDR146C | SWI5 | Swi5 |
| YDR113C | PDS1 | Pds1 |
| YAR019C | CDC15 | Cdc14 |

Shown are the systematic gene names, the standard names, and the corresponding network node names.

**Supplementary Table 5.** GRN used for hematopoietic stem cell dataset.

| Source | Target | Type |
|---|---|---|
| GATA2 | GATA2 | Activation |
| GATA2 | GATA1 | Activation |
| GATA1 | PU.1 | Inhibition |
| PU.1 | GATA1 | Inhibition |
| GATA1 | GATA1 | Activation |
| PU.1 | PU.1 | Activation |
| GATA1 | Fog-1 | Activation |
| PU.1 | Scl | Inhibition |
| GATA1 | Scl | Activation |
| GATA1 | c-Kit | Inhibition |
| Scl | c-Kit | Activation |
| Fog-1 | GATA2 | Inhibition |
| Fog-1 | c-Myb | Inhibition |
| GATA2 | PU.1 | Inhibition |
| PU.1 | GATA2 | Inhibition |
| GATA1 | c-Myb | Activation |
| GATA1 | GATA2 | Inhibition |
| GATA1 | Hbaa1 | Activation |
| c-Myb | GATA1 | Inhibition |
| PU.1 | CD11b | Activation |
| PU.1 | Fog-1 | Inhibition |

Original network topology was obtained from prior work by Mojtahedi et al, 2016, in PLoS Biology 14(12), referenced in the main text. We removed genes with low measured variance (Runx1, cJun, C/EBPa, EpoR, Eklf, Egr-2, Gfi-1, Fli-1).

**Supplementary Table 6.** Parameters and input gene expression space used for STICCC on synthetic and experimental circuits.

| Dataset | Type | Sampling Radius | Expression Space | Number of Models/Cells |
|---|---|---|---|---|
| REP | Simulated | 0.05 | All PCs | 10000 |
| CTS | | | | |
| iFFL | | | | |
| TS/REP | | | | |
| REP - time series | | 0.15 | | 2000 |
| REP - down-sampled | | 0.1 | | 200 |
| REP - dropout | | 0.1 | | 9994 |
| CTS - time series | | 0.15 | | 5000 |
| CTS - dropout | | 0.2 | | 2000 |
| CTS - signaling | | 0.05 | | 4377 |
| Cell cycle | Experimental | 0.15 | Top 10 PCs | 976 |
| HSC | | 0.2 | Top 5 PCs | 1600 |
| EMT | | 0.3 | Top 15 PCs | 3133 |
